# Distinct microbial communities are linked to organic matter properties in millimetre-sized soil aggregates

**DOI:** 10.1101/2024.08.01.606122

**Authors:** Eva Simon, Ksenia Guseva, Sean Darcy, Lauren Alteio, Petra Pjevac, Hannes Schmidt, Kian Jenab, Christian Ranits, Christina Kaiser

## Abstract

Soils provide essential ecosystem services and represent the most diverse habitat on Earth. It has been suggested that the presence of various physico-chemically heterogenous microhabitats supports the enormous diversity of microbial communities in soil. However, little is known about the relationship between microbial communities and their immediate environment at the micro- to millimetre-scale. In this study, we examined whether bacteria, archaea, and fungi organise into distinct communities in individual 2-millimetre-sized soil aggregates and compared them to communities of homogenized bulk soil samples. Furthermore, we investigated their relationship to their local environment by concomitantly determining microbial community structure and physico-chemical properties from the same individual aggregates. Aggregate-communities displayed exceptionally high beta-diversity, with 3-4 aggregates collectively capturing more diversity than their homogenized parent soil core. Up to 20-30% of ASVs (particularly rare ones) were unique to individual aggregates selected within a few centimetres. Aggregates and bulk soil samples showed partly different dominant phyla, indicating that taxa that are potentially driving biogeochemical processes at the small scale may not be recognized when analysing larger soil volumes. Microbial community composition and richness of individual aggregates were closely related to aggregate-specific carbon and nitrogen content, carbon stable-isotope composition, and soil moisture, indicating that aggregates provide a stable environment for sufficient time to allow co-development of communities and their environment. We conclude that the soil microbiome is a metacommunity of variable subcommunities. Our study highlights the necessity to study small, spatially coherent soil samples to better understand controls of community structure and community-mediated processes in soils.

## Introduction

Soil harbours a huge microbial diversity [1, 2], with one gram of soil containing tens of thousands of bacterial, archaeal, and fungal taxa [1, 3]. At the same time, soil is probably the most structurally intricate microbial environment on Earth, consisting of a complex three-dimensional architecture of mineral particles, organic material, and pore spaces [4]. This complexity leads to significant variability in environmental conditions at small scales, which give rise to a large variety of physically and chemically distinct microhabitats [5–9]. The overwhelming richness of the soil microbiome may thus be the result of an unseen high beta-diversity of communities at small scales [6, 9, 10]. Not much, however, is known about the small-scale structure of the soil microbiome.

Soil microbial communities regulate large-scale biogeochemical cycles by carrying out highly specialized metabolic functions and biochemical processes at the small scale in close interaction with their abiotic environment [3]. Although the composition and richness of the soil microbiome have been found to be related to soil biogeochemistry across different soils or treatments [3, 11–16], identifying a more general link between specific soil biogeochemical processes or environments and community structure remains a challenge [3, 6]. One reason for these difficulties might be the disparity of spatial scales between the size of microhabitats relevant to microbial communities and the size of studied soil samples. Bacterial interactions and dispersal are restricted to short distances as discontinuous water films within most soils limit bacterial movement and diffusion of metabolites [17–19]. Bacteria cluster into micro-scale patches throughout the soil pore space [5, 7, 17, 20–23] which may be linked to the heterogenous distribution of chemically diverse organic matter [17, 22, 24–26]. Fungi, in contrast, interact with larger soil volumes than bacteria and archaea, and are due to their hyphal lifestyle probably less dependent on small-scale environmental conditions. Hence, if we define microbial communities as “a set of potentially interacting species that co-occur in space and time” [10], bacterial and archaeal communities are likely confined to micro- or millimetres, whereas fungal communities may span larger spatial scales of unknown extent. Whether there is a relation between communities and their environment, can only be observed when investigated at its appropriate scale.

To date, both community structure and soil functions are mostly measured in ‘bulk soil samples’, that are aliquots of large soil volumes produced by homogenising one or more soil cores (typically covering hundreds to thousands of cubic centimetres) through sieving. Although this handling is useful for gaining information representative of a larger soil volume, it destroys information regarding the spatial distribution of microbial communities and their potential link with their in-situ environment [7]. Using homogenized bulk soil samples for studying relationships between community structure and soil properties implies that patterns observed at the larger scale reflect those at the small scale. However, whether this assumption is valid is not known.

Studying individual soil aggregates offers an interesting approach for gaining deeper insights into microbial community structure and its link to the surrounding environment at the millimetre-scale [18, 27–29]. Aggregates are micro- to millimetre-sized conglomerates of mineral particles and organic material which cohere after soil disintegration as bonds between their particles are stronger than bonds with surrounding particles. Aggregates are proposed to pose a scale that approximates connected bacterial communities [18, 27–29]. However, only very few studies have investigated bacterial communities in individual soil aggregates [30–32]. To our knowledge, until today, no study has concomitantly analysed community structure and physico-chemical properties of individual aggregates to investigate a potential link at this small scale.

Whether microbial communities are structured into distinct subcommunities at the small-scale depends on the temporal stability of their immediate environment. It has been proposed that soil aggregates, which are stable for weeks to months [33], provide a stable environment for enough time to allow evolution and assembly of distinct microbial communities [34]. If this is the case, we expect that microbial richness and community composition are linked to chemical properties (such as carbon or nitrogen content) at the scale of individual aggregates.

In this study, we address two main questions: (i) Do bacteria, archaea, and fungi organise into distinct communities at the scale of individual aggregates that are qualitatively different from those observed in bulk soil samples? Answering this question will allow us to evaluate the necessity to sample smaller scales to gain insights into structure-function links. If aggregates host distinct communities, (ii) what are the main drivers of microbial community structure of individual aggregates? Is community structure influenced by the physico-chemical environment and/or the spatial distance between communities? To answer these questions, we collected 190 2-millimetre-sized individual aggregates from 40 soil cores (“bulk soil samples”) of known geographical location in two soil depths across a forest site. We measured bacterial, archaeal, and fungal community composition, richness and density, and carbon and nitrogen content, isotopic composition of carbon and nitrogen, C-to-N ratio, and soil water content concomitantly from the same aggregates. We (i) compared community properties between aggregates and bulk soil samples and (ii) explored the link between communities and their physico-chemical environment across individual aggregates.

## Material and Methods

### Study site, sample collection and processing

We collected soil cores from a temperate beech-dominated forest in Klausen-Leopoldsdorf, Austria (48.11925 N, 16.04343 E, 580 m altitude) in autumn 2020. The soil type is classified as a dystric cambisol over sandstone with a loam-loamy clay texture [35–37]. We chose 10 sampling points across a 11 x 65 meters-large southwest facing-slope and recorded their relative position to each other. Always two sampling points lay approximately one meter apart within the same plot (5 x 5 meters). At each sampling point, we took two soil cores, one from 0-5 centimetres soil depth and another one from 15-20 centimetres. From each core, we subsampled two soil cores (35 cm^3^ volume, 3 cm diameter, 5 cm height) approximately one centimetre apart, which we term “parent soil cores*”.* From each parent soil core (n=40), we hand-picked individual approximately 2-millimetre-sized soil clumps (6.7 ± 2.2 mg dry soil), referred to as “aggregates*”* and sieved remaining soil to two millimetres, producing “bulk soil samples” (see Fig. S1 for a detailed sampling scheme). For hand-picking individual soil aggregates we transferred a small amount of each soil sample in the centre of a petri-dish which contained a ring of moist filter paper. The petri-dish was always closed with a lid which was only opened when an aggregate was selected. We then selected 5 aggregates, one after another, from each sample. Each selected aggregate was placed in a tared metal tin with the help of a fine tweezer and weighed on a fine mass balance (precision: 0.000001 g) to determine its fresh weight. After weighing, the aggregate was transferred into a safe-lock tube which we closed immediately. Aggregates in individual tubes were stored at -80°C. The whole procedure of picking an aggregate over weighing it until closing the tube took only a few minutes. With this approach we aimed to ensure as little drying of soil aggregates during handling as possible (for more details see supplementary Material and Methods). Still, we cannot fully rule out that environmental conditions of aggregates, especially at their surface, might have changed upon sampling due to exposure to air. However, we do not expect these changes to have affected measurements of microbial community composition and richness as turnover times of microorganisms lie in the range of days to weeks and not minutes to hours [38–40].

We freeze-dried bulk soil samples and individual aggregates (for 48 h) which had been stored at -80°C. We weighed freeze-dried, still intact individual aggregates before carefully homogenising them by hand using a stainless-steel ball tool to produce homogeneous samples for further analyses.

Based on preliminary assessment of homogenisation of aggregates, we were confident that analysed aggregate-aliquots were representative of the whole aggregate concerning community and elemental composition (Fig. S2).

### DNA extraction and sequencing

DNA was extracted from freeze-dried and homogenised aliquots of aggregates (n=190, 3.8 ±1.8 mg dry soil) and bulk soil samples (n=40, 197.5 ± 35 mg dry soil) with the Qiagen PowerSoil Pro Kit (Qiagen, Valencia, CA, USA). We followed the manufactureŕs protocol, except that we replaced the 10-minute vortexing step with a bead-beating step (30 seconds) on a FastPrep-24 Classic Bead Beater (MP Biomedicals, Germany) for improved cell lysis.

We amplified the V4 region of the 16S rRNA genes targeting bacteria and archaea, and the fungal internal transcribed spacer region 2 (ITS2) of the nuclear rRNA operon, following a two-step PCR barcoding approach [41]. For the amplification of the 16S rRNA genes, we used primers 515F [42] and 806R [43]. For the ITS2 region, we used a nested-like approach [44] whereby we first generated a PCR product using primers ITSOF-T [45, 46] and ITS4 [47], and then used it as a template for a second PCR using primers gITS7 [48] and ITS4 (see PCR cycling conditions in Table S1). The final product was subsequently barcoded [42]. Amplicons were sequenced on a MiSeq platform (Illumina, V3 chemistry, 600 cycles) at the Joint Microbiome Facility of the Medical University of Vienna and the University of Vienna.

We extracted amplicon pools from raw sequencing data using the FASTQ workflow in BaseSpace (Illumina, default settings), removed PhiX contamination using BBDuk (BBtools, [49]), and demultiplexed data using the package demultiplex (Laros JFJ, github.com/jfjlaros/demultiplex) allowing one mismatch for barcodes, and two for linkers and primers. We inferred amplicon sequence variants (ASVs) with the DADA2 package (version 1.18.0 [50]) with default settings. Finally, 16S rRNA genes ASVs were classified against the SILVA database (SSU RefNR 99, release 138.1) using the classifier implemented in DADA2. Fungal ITS2 ASVs were classified against the UNITE database (Version 04.02.2020 [51]).

### Quantification of bacterial, archaeal, and fungal abundances

The concentration of DNA extracts was measured with a HS dsDNA assay following the manufacturer’s protocol on a Qubit 4.0 fluorometer (Thermo Fisher Inc, Waltham, MA, USA). We assessed concentration of bacterial and archaeal 16S rRNA gene copies and fungal ITS regions of DNA extracts with the QX200 Droplet Digital PCR (Bio-Rad). DNA extracts were diluted to 0.1 ng (16S rRNA genes) and 0.5 ng (ITS1 region) of total DNA per reaction to optimize the separation of negative and positive droplets. See Table S2 for primers and PCR cycling conditions. Droplet readings (>10000 droplets, ≥250 positive droplets, ≥250 negative droplets) were analysed using the QuantaSoft software (Bio-Rad).

### Soil water content, carbon and nitrogen content, **δ**^13^C, and **δ**^15^N

We assessed gravimetric soil water content of individual samples by subtracting weight of freeze-dried intact aggregates (n=137) and bulk soil samples (n=40) from their fresh weight assessed before 48 hours freeze-drying. For the aggregates, we weighed each intact sample twice: i) when they were field-moist directly after they were hand-picked, and ii) after they had been freeze-dried.

We determined carbon and nitrogen content, and isotopic ratio of stable carbon and nitrogen isotopes of aliquots of freeze-dried, homogenised aggregates (0-5 cm: n=92, 0.5 ± 0.04 mg freeze-dried soil, 15-20 cm: n=98, 0.6 ± 0.03 mg dry soil) and freeze-dried, milled bulk soil samples (0-5 cm: n=20, 8.3 ± 0.4 mg dry soil, 15-20 cm: n=20, 12.1 ± 0.5 mg dry soil) via elemental analyser isotope ratio mass spectrometry with an elemental analyser (EA IsoLink CN, Thermo Scientific) coupled to an isotope ratio mass spectrometer (Delta V Advantage Isotope Ratio MS Hi Pan CNOS, Thermo Scientific) via a ConFlo IV interface (Thermo Scientific).

### Data analyses

Data analyses were performed in R 4.2.0 version [52]. We used packages TreeSummarizedExperiment [53], phyloseq [54], and vegan [55] for sequencing data analyses. Graphs were generated with ggplot2 [56] and edited in Inkscape (v.0.92.4, [57]).

Community composition analyses were based on non-rarefied datasets (aggregates: 0-5 cm: n=92, 15-20 cm: n=98, bulk soil samples: 0-5 cm: n=20, 15-20 cm: n=20). Analyses examining microbial richness (number of ASVs) were based on rarefied datasets, including 186 (16S rRNA genes) and 183 (ITS2) aggregates and 39 (16S rRNA genes) and 38 (ITS2) bulk soil samples after rarefying samples to 8,185 (16S rRNA genes) or 8,195 (ITS2) reads.

We generated mean species accumulation curves from rarefied count tables for (i) aggregates and bulk soil samples to show the variability of samples; and for (ii) aggregates from the same parent core to visualise the overlap of ASVs between spatially proximate aggregates.

We visualised beta-diversity of bacterial and fungal communities of aggregates and bulk soil samples with principal coordinate analysis (PCoA) plots. We performed PCoAs on Aitchison distances, which are Euclidean distances calculated from centered-log-ratio transformed counts matrix [58] and are appropriate to address the compositional nature of amplicon sequencing data [59]. We analysed the relationship of community dissimilarity (Aitchison distance) and spatial distance between samples (Euclidian distance). For plotting, we discretised community dissimilarity into spatial distance categories based on the approximate distance between sample pairs.

To assess the similarity of communities among aggregates from the same parent soil core, we determined (i) how many ASVs of an individual community were exclusively present in one out of four aggregates or shared with three other aggregates, and (ii) which fraction of a community exclusive and shared ASVs made up.

To assess if aggregates and bulk soil samples suggest the same phyla to dominate the soil microbiome, we compared phylum composition of 100 abundance categories, which contained different numbers of the most abundant ASVs (per sample) across all aggregates and bulk soil samples. In each sample, we ranked ASVs according to their relative abundance and selected the 100 most abundant ASVs as they are often assumed to be especially important for community functioning. Based on their ranks we binned ASVs from all samples into cumulative abundance categories (category 1 to 100) across aggregates and bulk soil samples divided by soil depth.

We performed variation partitioning to quantify the relative importance of three predictor sets: (i) parent soil core belonging, (ii) spatial distribution, and (iii) elemental composition of aggregates for microbial community composition and richness in both soil depths. Soil core belonging was included as a dummy variable. We calculated spatial distribution as principal coordinates of neighbour matrices (PCNM). We performed forward selection for PCNM axes and elemental composition variables (C and N content, C-to-N ratio, δ^13^C, and δ^15^N), which were centred and scaled to guarantee their equal weight in forward selection of variables. Which variables were used per predictor set ii and iii depended on investigated response variable and soil depth.

We used linear regression models to elucidate the relationship between microbial community composition and z-transformed C and N content, C-to-N ratio, δ^13^C, δ^15^N, and soil water content at the aggregate-scale. We only consider aggregates from the same parent soil core to minimise the veiling effect of heterogeneity at the meter-scale.

To examine the relationship between richness and C, N content, C-to-N ratio, δ^13^C, δ^15^N, soil water content, and microbial density of individual aggregates, we performed linear and quadratic polynomial regression models. We present the model with better fit based on ANOVA and AIC.

## Results

### Microbial communities were more variable across aggregates than bulk soil samples

Both bulk soil samples and aggregates hosted unique microbial communities (Fig. 1, Fig. 2). Species accumulation curves of bulk soil samples reached a plateau with fewer samples than curves of aggregates (Fig. 1). The marked variability of microbial communities across aggregates is also visible in ordination plots (Fig. 2). Comparing the degree of community dissimilarity between aggregates and bulk soil samples at particular spatial distance, we observed that microbial communities in aggregates exhibited a higher dissimilarity than bulk soil samples, in particular at close distances of a few centimetres (Fig. 3).

**Fig. 1.**
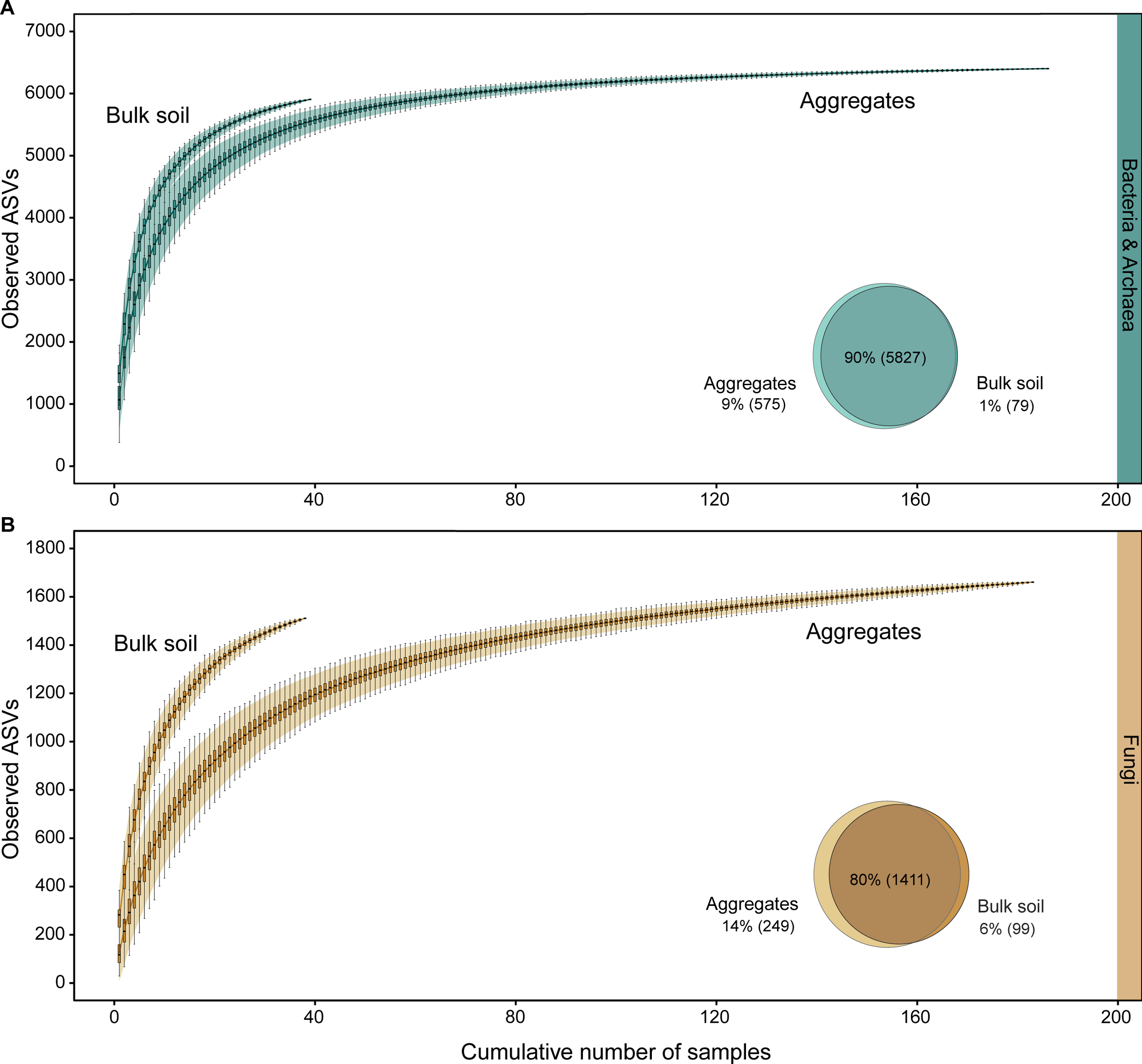
Aggregates and bulk soil samples, respectively, partly varied in microbial ASVs. Mean species accumulation curves generated from 1,000 random permutations display the cumulative number of observed bacterial and archaeal, and fungal ASVs in relation to sampling effort (cumulative number of aggregates and bulk soil samples, respectively). Venn diagrams depict shared and exclusive ASVs between aggregates and bulk soil samples. Species accumulation curves and venn diagram for (**A**) bacteria and archaea (green) and (**B**) fungi (orange) are calculated from rarefied ASV count tables (correcting for the effect of varying sequencing depths of samples) for bulk soil samples (Bulk soil, 16S rRNA genes: n=39, ITS2: n=38) and aggregates (Aggregates, 16S rRNA genes: n=186, ITS2: n=183). Species accumulation curves are produced by randomly adding samples to the accumulation curve and plotting the mean of permutations. The shaded area around curves represents 95% confidence interval. Individual boxplots display median number of unique ASVs in a cumulative number of samples. Venn diagrams in the lower right corner of plot windows indicate the percentage of bacterial and archaeal or fungal ASVs which were detected in both, aggregates and bulk soil samples or detected exclusively in one sample type.

**Fig. 2.**
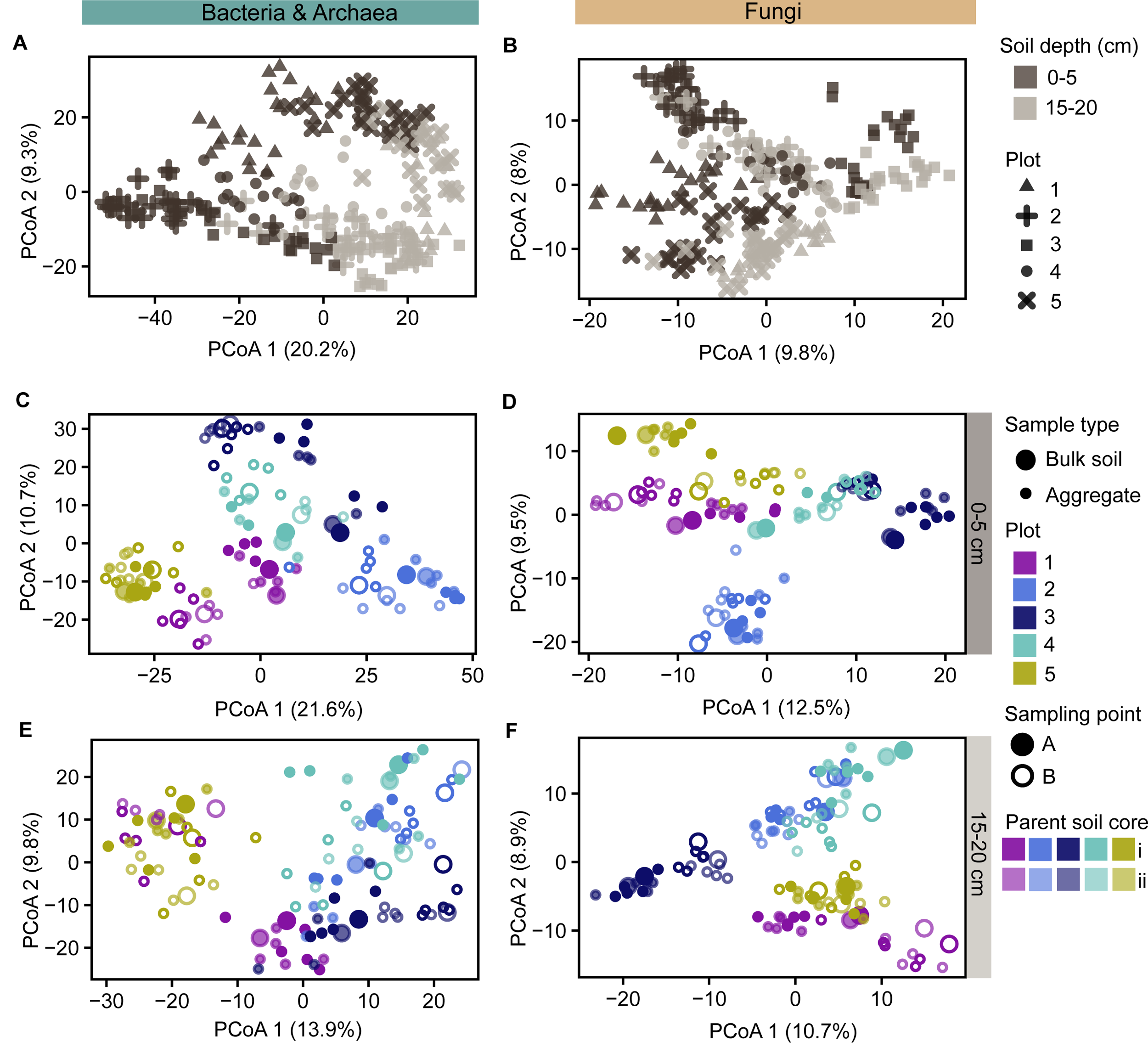
Variability in microbial community composition was associated with soil depth and sampling location. Principal coordinate analysis (PCoA) plots depict variation in bacterial and archaeal and fungal community composition across (**A, B**) aggregates (n=190) and bulk soil samples (n=40) of both soil depths, (**C**, **D**) of 0-5 centimetres and (**E**, **F**) 15-20 centimetres soil depth. PCoA plots were generated using Aitchison distances as the metric for microbial community similarity which were calculated from non-rarefied ASV count tables. Each point represents a sample (aggregates depicted by smaller points, bulk soil samples by larger points). The closer samples are ordinated, the more similar their community composition. Variation in microbial community similarity explained by the first and second PCoA axis is given in parentheses. (**A, B**) Symbol colour depicts soil depth (dark brown for 0-5 centimetres, light brown for 15-20 centimetres soil depth), and symbol shape indicates the plot (1-5) from which parent soil cores were sampled. (**C, D, E, F**) Unique combinations of symbol colour, shape and opacity indicate parent soil core belonging of samples: Symbol colour represents the plot (1-5) from which parent soil cores were sampled, symbol shape indicates the sampling points of a plot which samples were taken (two per plot, which were taken one meter apart from each other), and opacity level depicts from which parent soil core per sampling point samples were derived (two parent soil cores per sampling point, few centimetres apart from each other). For further information on the sampling scheme see Fig. S1.

**Fig. 3.**
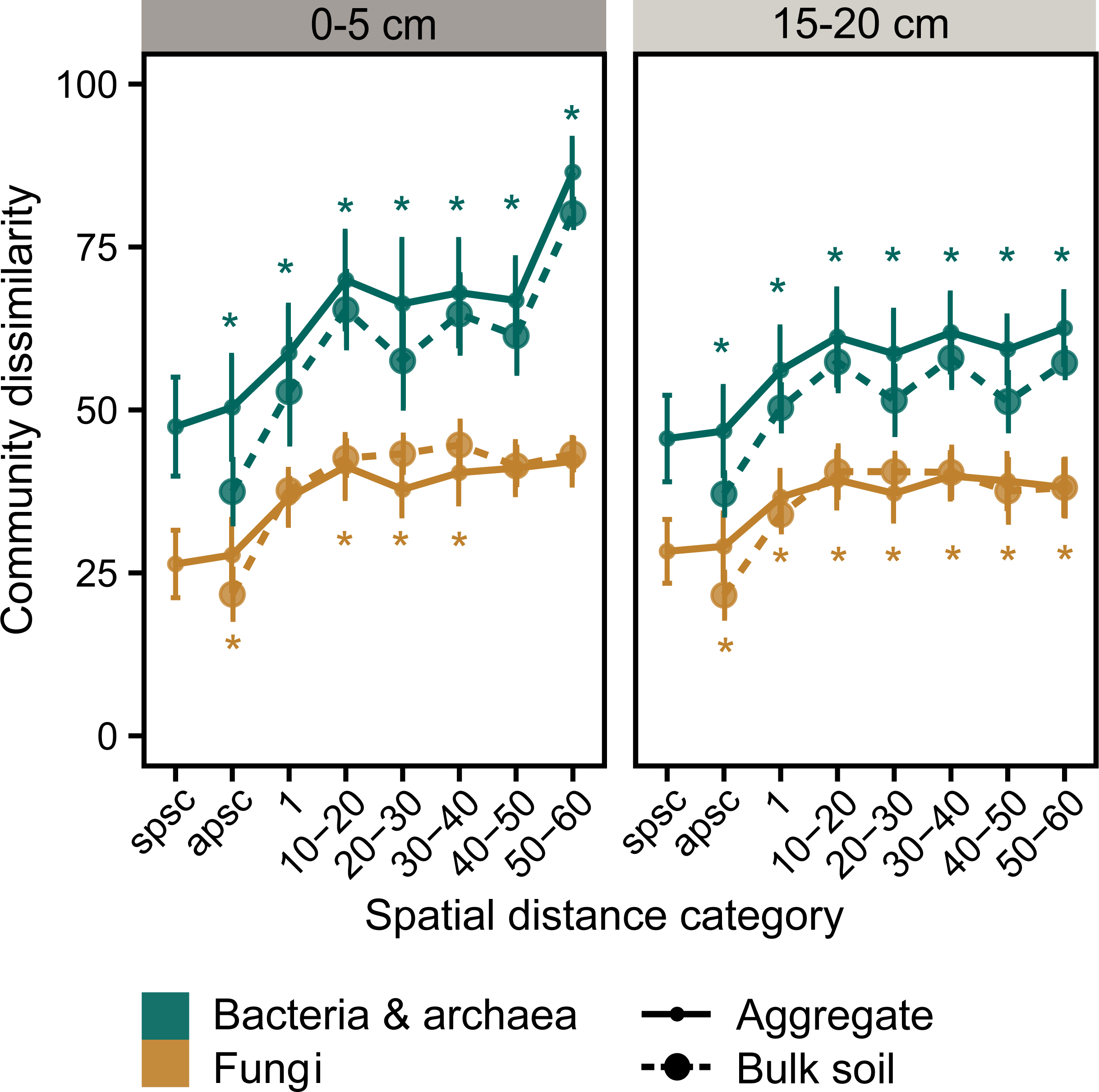
Community composition of bacteria and archaea were at the same spatial distance more dissimilar from each other in aggregates than in bulk soil samples. Spatially most proximate samples were most similar in community composition. Plots display the relationship of bacterial and archaeal (green) and fungal (orange) community composition and spatial distance between samples in 0-5 centimetres and 15-20 centimetres soil depth. Community dissimilarity is calculated as Aitchison distances from the non-rarefied datasets, spatial distance between sample pairs is calculated as Euclidean distances. Community dissimilarities are grouped into spatial distance categories based on the relative spatial distance between samples: (i) milli- to centimetres apart as samples were from the same parent soil core (spsc), (ii) centimetres apart as samples were from adjacent parent soil cores (apsc), (iii) approximately one meter apart (1), (iv-viii) approximately 10-20, 20-30, 30-40, 40-50 or 50-60 meters apart. Data points represent average community dissimilarity (depicted as medians) of distinct spatial distance categories (x-axis). Error bars indicate the standard deviation of the median. Small points and solid line type represent aggregates, large points and dashed line type bulk soil samples. Different numbers of pairwise comparisons underlie individual data points (aggregates: n=152-950, bulk soil samples: n= 9-40), for a complete list see Table S3. Green asterisks indicate significant difference (*P* value< 0.05) in community dissimilarity between aggregates and bulk soil samples for bacterial and archaeal communities and orange asterisks for fungal communities. Statistical significances were determined by Wilcoxon test between medians.

Although aggregates from the same parent soil core were overall compositionally most similar (Fig. 3), community composition of some aggregates was more similar to aggregates from other sampling locations or bulk soil samples than to aggregates of the same parent soil core (Fig. 2). This underpins the high variability and heterogeneity of aggregate-associated microbial communities and suggests that they were, at that small scale, not solely determined by sampling location.

### Most-similar aggregates differed in low abundant ASVs

Although aggregates from the same parent soil core shared the most abundant ASVs, they considerably differed in low abundant ASVs (Fig. 4C-F). On average, 37% and 31% of bacterial and archaeal, and fungal ASVs were shared among four aggregates belonging to the same parent soil core. These shared ASVs accounted for more than three-quarters of total ASVs in each aggregate (78% bacteria and 86% fungi) (Fig. 4E, 4F). However, 20% and 27% of bacterial and archaeal, and fungal ASVs were unique to one of the four aggregates (Fig. 4C, 4D), exemplifying the high heterogeneity of microbial communities at the millimetre-scale. Unique ASVs made up for only 4% and 3% of total abundances of an average bacterial and archaeal, and fungal community (Fig. 4E, 4F). Overall, we captured bacterial and archaeal, and especially fungal communities of aggregates more completely than of bulk soil samples (Fig. S3).

**Fig. 4.**
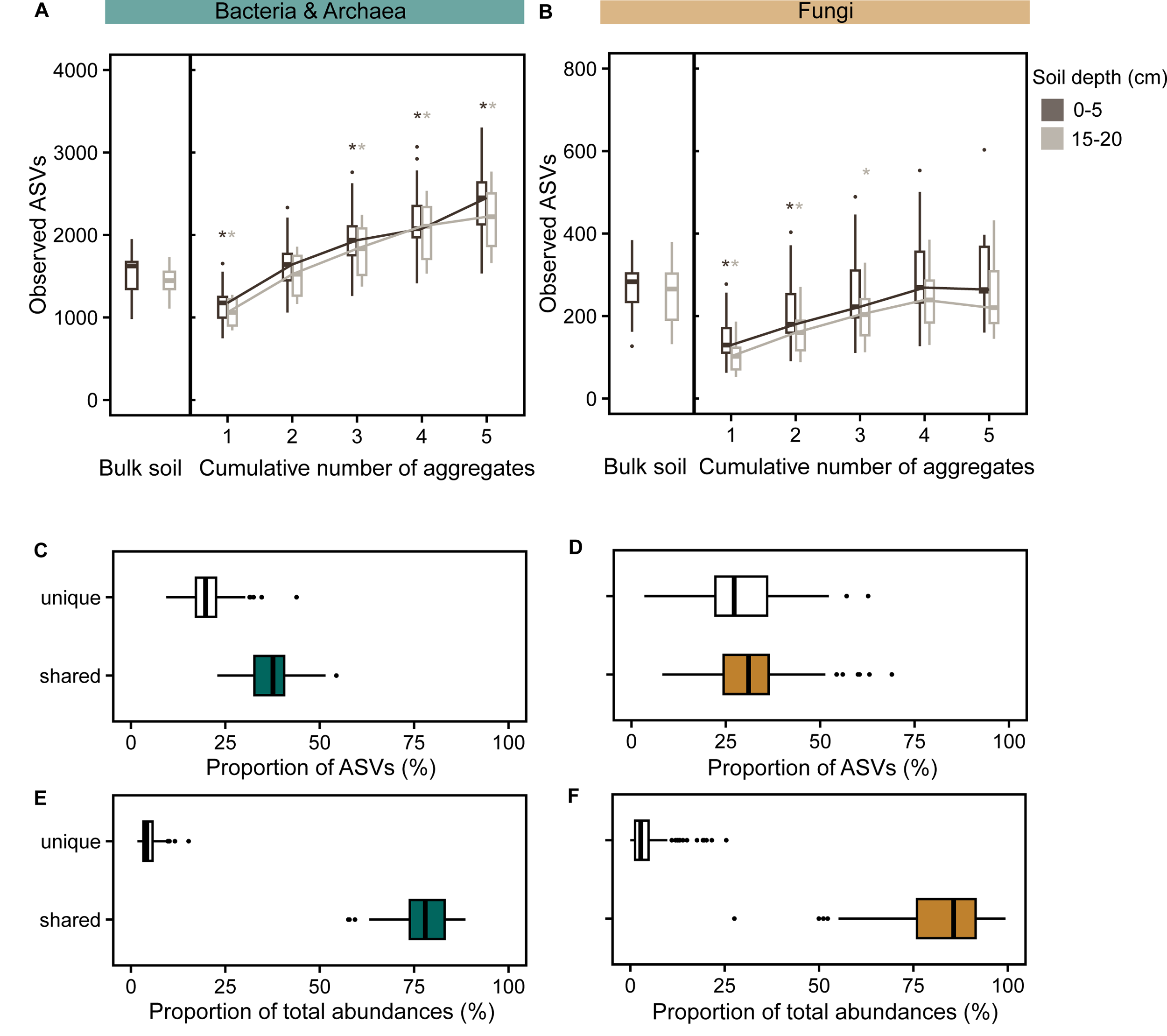
A few aggregates captured a higher diversity compared to their homogenised parent soil core. Plots are based on rarefied datasets. (**A, B**) The two boxplots to the very left of each plot illustrate average richness of (**A**) bacterial and archaeal and (**B**) fungal ASVs in rarefied bulk soil samples in 0-5 (dark brown) and 15-20 (light brown) centimetres soil depth. The five boxplots to the right represent the number of (**A**) bacterial and archaeal, and (**B**) fungal ASVs in one to five aggregates from the same parent soil core cumulatively in 0-5 (dark brown) and 15-20 (light brown) centimetres soil depth. They are based on individual species accumulation curves of aggregates (n=3-5) from the same parent soil core which were generated with 100 random permutations, only considering parent soil cores with at least three available aggregates (16S rRNA genes: n=38, ITS2: n=38). Asterisks above boxes indicating richness of aggregate(s) indicate significant difference (*P* value<0.05) between medians of bulk soil samples and the (cumulative number of) aggregate(s), which were calculated using Wilcoxon tests. Colour of asterisks depicts soil depth significances refer to. (**C, D**) Plots C and D show the proportion of (**C**) bacterial and archaeal and (**D**) fungal ASVs detected either exclusively (unique, white fill) in a single aggregate-community or found in four aggregates of the same parent soil core (shared, green or orange fill). (**E, F**) Plots E and F depict the proportion of relative abundances of an individual aggregate-community composed of unique (unique, white fill) and shared (shared, green or orange fill) (**E**) bacterial and archaeal and (**F**) fungal ASVs. Boxplots in plot C, D, E and F show the median and interquartile range (25-75% of data) across parent soil cores (16S rRNA genes: n=36; ITS2: n=38 parent soil cores).

### Aggregates revealed unexplored bacterial and archaeal richness

Aggregates and bulk soil samples depicted an equal cumulative richness of the soil microbiome at our field site (Fig. 1). However, when we examined bacterial and archaeal richness of individual parent soil cores, already three 2-millimetre-sized aggregates revealed a larger number of bacterial and archaeal ASVs (0-5 cm: median=1936, 15-20 cm: median=1832) compared to one bulk soil sample (0-5 cm: median=1623, 15-20 cm: median=1447). In the case of fungal richness, one bulk soil sample comprised slightly more fungal ASVs (0-5 cm: median=283, 15-20 cm: median=265) than five aggregates cumulatively (0-5 cm: median=264, 15-20 cm: median=220) (Fig. 4B).

### Aggregates and bulk soil samples differed in phylum composition of their most abundant bacterial and archaeal taxa

We found that above 90% (bacteria and archaea) and 80% (fungi) of total ASVs in our datasets were detected in both sample types (Fig. 1, venn diagrams). Some taxa of bacteria and archaea (9%) and fungi (14%) were exclusively observed in aggregates, and few taxa of fungi (6%) only in bulk soil.

When considering only most abundant ASVs in samples, aggregates and bulk soil samples partly depicted different phyla to dominate the soil microbiome, suggesting that we may over- or underestimate the role of some phyla in biogeochemical processes if we only examine bulk soil samples. Differences were more pronounced for bacteria and archaea (Fig. 5) than fungi (Fig. S6). For instance, *Actinobacteriota* were amongst the topmost abundant phyla in aggregates but not in bulk soil samples (Fig. 5). In contrast, members of *Crenarcheota* and *Planctomycetota* were detected amongst the 100 most abundant ASVs in bulk soil samples but not in aggregates.

**Fig. 5.**
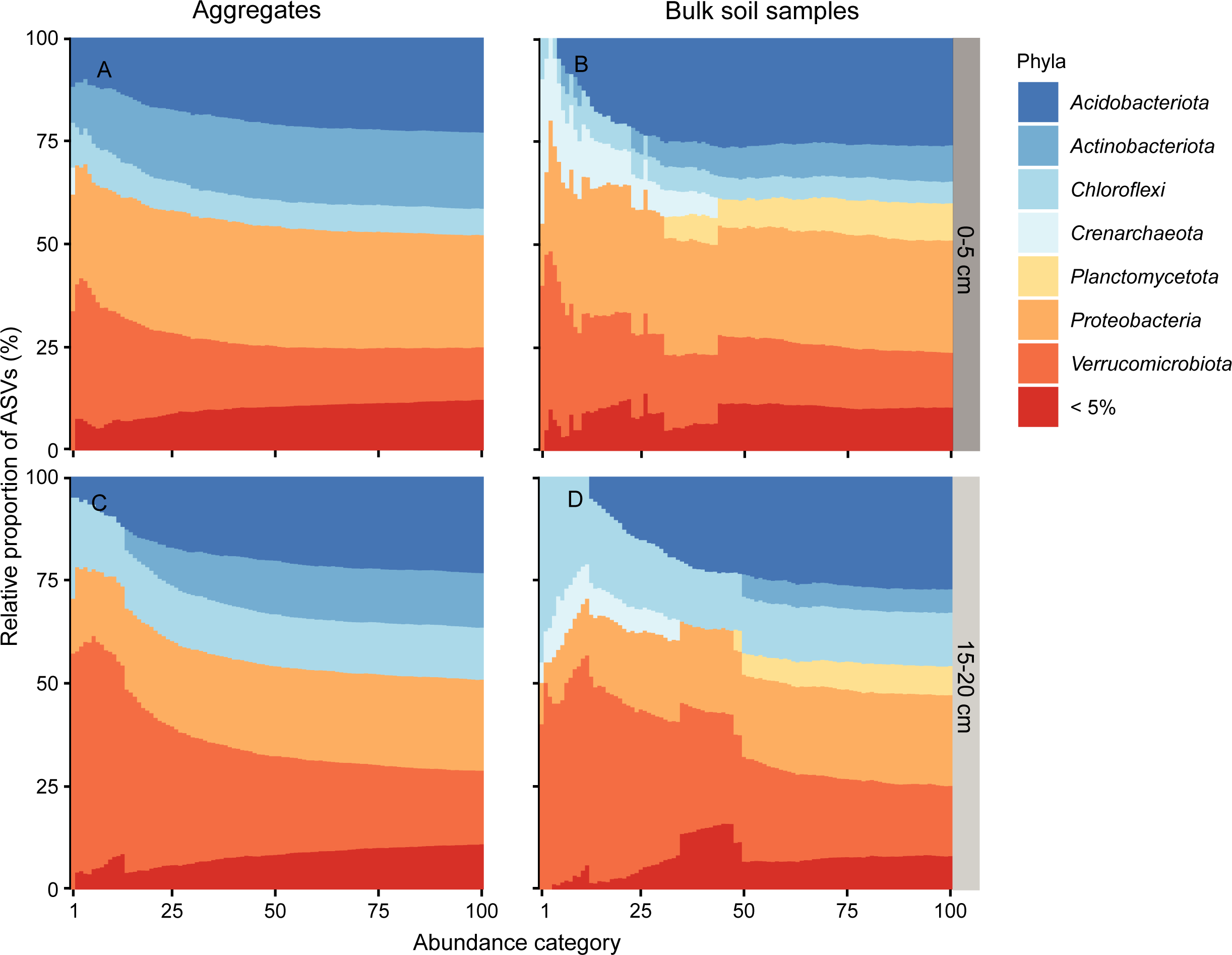
Aggregates and bulk soil samples differed in phylum composition of their most abundant bacterial and archaeal taxa. Stacked bar plots illustrate the phylum composition of 100 abundance categories for (**A, C**) aggregates (0-5 cm: n=92,15-20 cm: n=98) and (**B, D**) bulk soil samples (0-5 cm: n=20, 15-20 cm: n=20) in 0-5 and 15-20 centimetres soil depth. Each stacked bar represents one abundance category (x-axis) and displays the proportions of ASVs belonging to different phyla (y-axis). Abundance categories comprise different numbers of ASVs. Which ASVs are included depends on their ranks which were assigned to ASVs in each sample individually based on their relative abundance. The number of abundance category (1 to 100) indicates how many of the most abundant ASVs (per sample) were included. For instance, abundance category 25 encompasses the 25 most abundant ASVs of all samples. Phyla comprising more than 5% of all ASVs per abundance category are depicted in different colours, whereas phyla constituting less than 5% of ASVs are grouped into the phylum category “<5%”. The analysis is based on non-rarefied datasets.

### Bacterial and archaeal aggregate-communities were strongly influenced by soil depth, whereas fungal communities were more influenced by horizontal sampling location

We observed that microbial communities differed between the topsoil (0-5 cm) and the deeper soil layer (15-20 cm) (Fig. 2). The difference was especially pronounced for bacteria and archaea, where we see a separation of bacterial communities by soil depth along the first principal component (Fig. 2A). Bacterial and archaeal, and, to a lesser extent, fungal communities were overall more variable in 0-5-centimetre than 15-20-centimetre soil depth (Fig. S5).

Horizontal sampling location had a strong effect on bacterial and archaeal, and fungal community composition (Fig. 2). Aggregates from the same and adjacent parent soil cores were on average most similar in composition (Fig. 3). Bacterial and archaeal community composition was linked to sampling location stronger in the topsoil than in the deeper soil layer, in which composition was only weakly associated with sampling location (Fig. 2A). Contrastingly, fungal communities were equally affected by sampling location in both depths (Fig. 2).

Between millimetres-to 10-20-meters-distance, community dissimilarity increased with spatial distance (Fig. 3). Beyond 10-20 meters the degree of similarity persisted irrespective across distances between samples. Only in topsoil, bacterial and archaeal community dissimilarity sharply increased between 40-50- and 50-60-meters distance (Fig. 3A).

### Variability of community composition was overall best explained by sampling location, but within soil core, also associated with carbon and nitrogen content

Parent soil core belonging explained by far the largest proportion of variation in microbial community composition of aggregates (Fig. 6A, 6B). Parent soil core belonging and spatial distribution of aggregates jointly accounted for about 20% of variation, indicating covariance of the two predictor sets. Individually, parent soil core belonging explained about 10% of variation (bacteria and archaea: 12% and 8%, fungi: 15% and 13%). Parent soil core belonging explained 12% (bacteria and archaea) and 8% (fungi) more variation in 0-5 centimetres than15-20-centimetres soil depth.

**Fig. 6.**
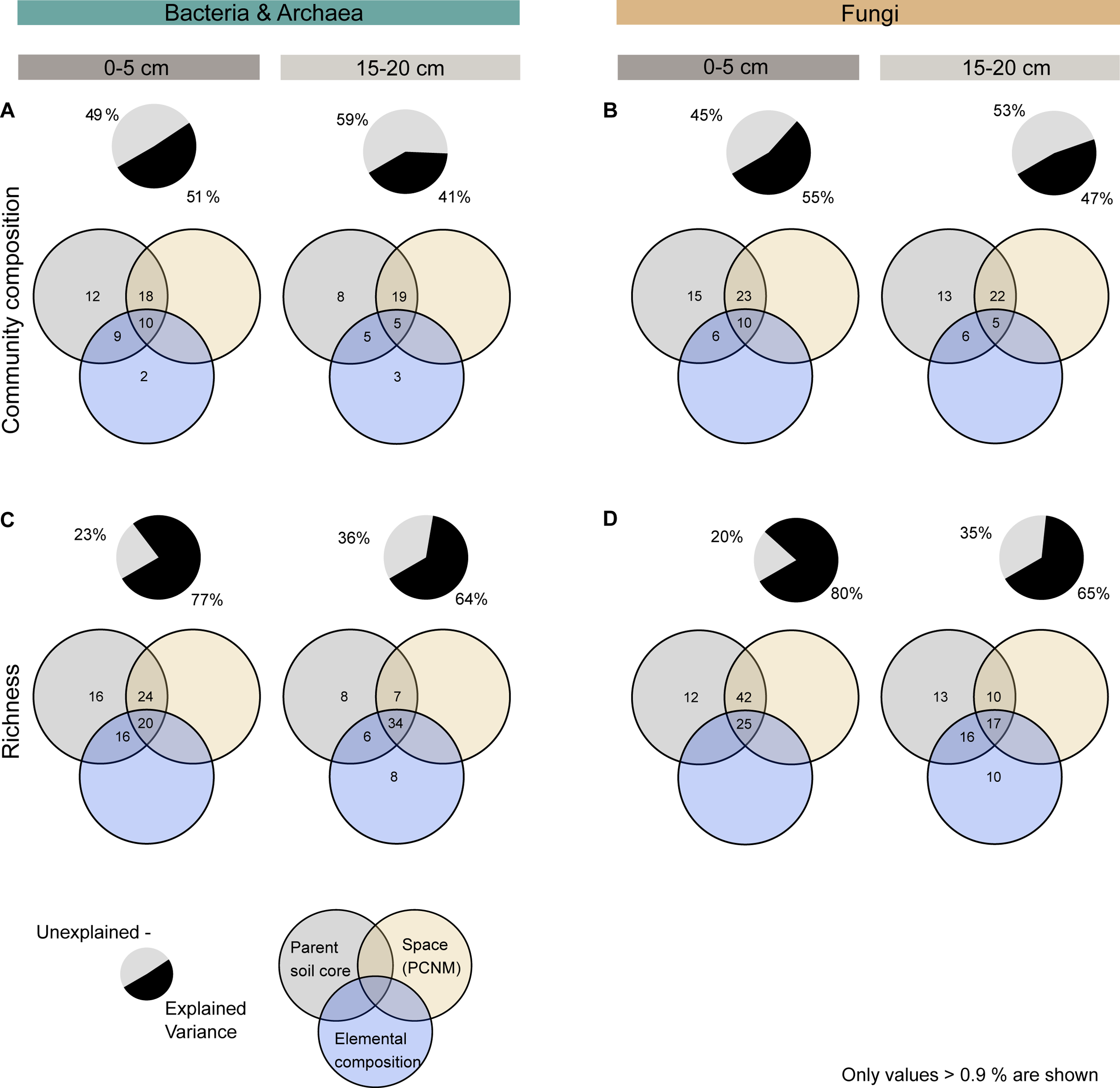
Although parent soil core belonging explained the largest proportion of variation in microbial community composition and richness of aggregates, elemental composition also played a role. Venn diagrams visualise the results of variation partitioning, illustrating the proportion (%) of variation in bacterial and archaeal, and fungal (**A, B**) community composition and (**C, D**) richness explained by one, two or three of the three predictor sets: (i) parent soil core belonging of aggregates (parent soil cores, grey), (ii) spatial distribution of aggregates (space, yellow), and (iii) aggregate-specific elemental composition of samples (elemental composition, blue) in soil depths 0-5 (left, n=80) and 15-20 (right, n=87) centimetres. Non-overlapping parts of circles depict the variation explained by an individual predictor set, whereas intersections of circles give variation explained by combinations of two or three predictor sets. Variation partitioning was performed on rarefied datasets. Here, we only display explained variation above >0.9%. Pie charts illustrate the proportion (%) of variation explained by the three predictor sets (black) and the remaining unexplained variation (light grey).

To filter out the strong effect of parent soil core belonging, we assessed the relationship between community dissimilarity and dissimilarity in environmental conditions among aggregates of the same parent soil cores. Our results demonstrated that the more similar C and N content of aggregates was, the more alike was their community composition (Fig. 7). Overall, community composition and C and N content were more strongly linked for bacteria and archaea than fungi. This was especially true for the deeper soil layer, where C and N content explained up to 26% of the total variability in bacterial community composition (R^2^ = 0.26, p<0.001). In contrast, δ^13^C (for bacteria, archaea, and fungi) and δ^15^N (for bacteria and archaea) were associated with community composition more strongly in the upper than the lower soil layer.

**Fig. 7.**
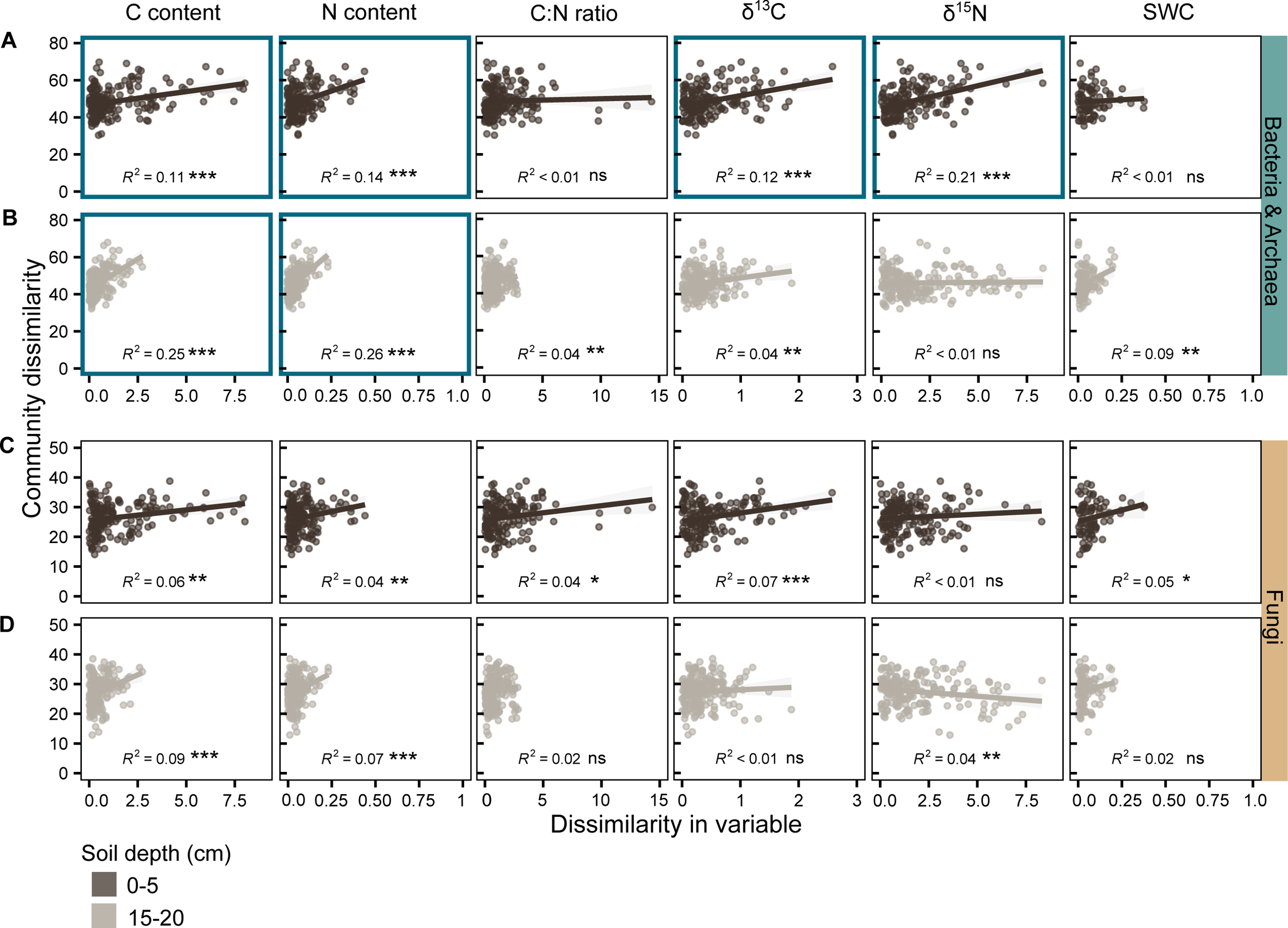
Bacterial and archaeal community composition was linked to C and N content and isotopic signatures of C and N in aggregates from the same parent soil core. Linear regression plots illustrate the relationships between dissimilarity of (**A, B**) bacterial and archaeal and (**C, D**) fungal community composition (Aitchison distances) and dissimilarity (Euclidean distances) of z-standardised carbon content (C content), nitrogen content (N content), carbon-to-nitrogen ratio (C:N ratio), δ^13^C (δ^13^C), δ^15^N (δ^15^N) values and soil water content (SWC) of aggregates at soil depths of 0-5 (dark brown) and 15-20 (light brown) centimetres. Dissimilarities were calculated for aggregates from parent soil cores with at least three available aggregates. Data points represent pairwise community dissimilarities (y-axis) and environmental variables (x-axis) of aggregate pairs from the same parent soil core. Lines depict calculated regression lines, with shaded areas around them indicating the 95% confident interval for the regressions. We print goodness-of-fit measure (*R*^2^, model accuracy) of the regression models, with asterisks denoting significance levels (*** *P* value≤0.001, ** *P* value≤0.01, * *P* value≤0.05). Highly significant correlations with a *R*^2^>0.1 are highlighted with a bold, coloured border around plots.

### Bacterial, archaeal, and fungal richness of aggregates was strongly linked to aggregate-specific C and N content, **δ**^13^C, **δ**^15^N, soil water content, and microbial density

As for community composition, parent soil core belonging explained the largest proportion of variation in fungal and bacterial richness in the upper soil layer. However, in the deeper soil layer, the relative importance of elemental composition was equal to sampling location (Fig. 6C, 6D).

Bacterial, archaeal, and fungal ASV richness were strongly correlated with aggregate-specific C and N content, δ^13^C and δ^15^N, and soil water content (Fig. 8). The nature of relationships between richness and environmental variables and microbial density partly differed between soil depths. In the topsoil, richness increased with C and N content until a maximum value beyond which it decreased again. Whereas, in the lower soil layer, richness showed a monotonic increase with growing contents. C-to-N ratio was positively related with bacterial and fungal richness only in the lower soil layer. In contrast, the more ^13^C-enriched aggregates were, the less ASV-rich they were, in both soil layers. Also, an increase in δ^15^N was associated with a decrease in bacterial, archaeal, and fungal richness. Furthermore, the higher the soil water content of an aggregate was, the more ASVs were observed. The relationship of gene copy numbers and richness followed unimodal shape in 0-5 centimetres for bacteria and archaea (Fig. 8E) but was linear in the deeper soil layer (Fig. 8F).

**Fig. 8.**
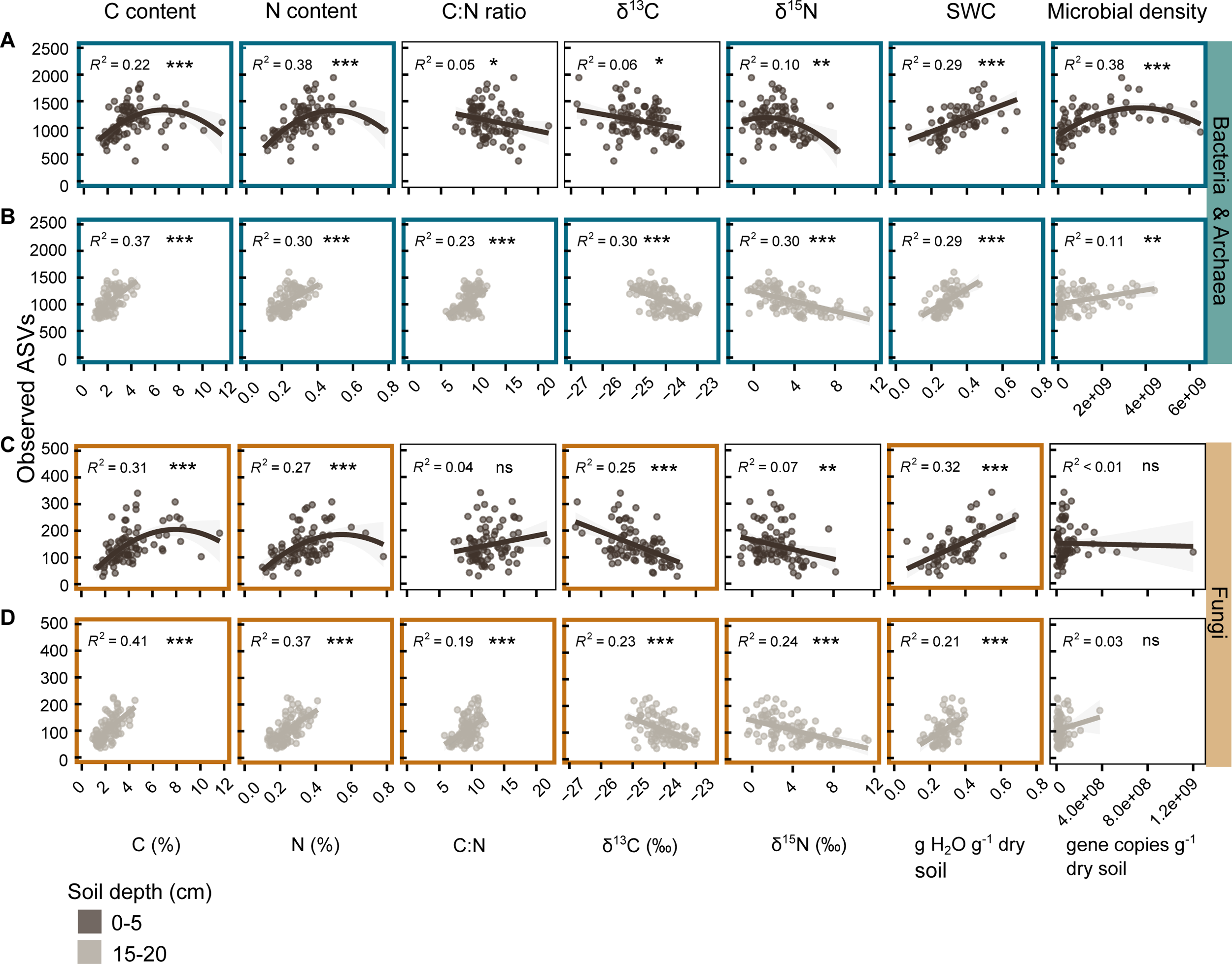
Microbial richness was strongly associated with C and N content and isotopic signatures of C and N, soil moisture and microbial density. Second-order polynomial and linear regression plots illustrate the relationships between the number of rarefied (**A, B**) bacterial and archaeal, and (**C, D**) fungal ASVs and carbon content (C (%)), nitrogen content (N (%)), carbon-to-nitrogen ratio (C:N), δ^13^C (δ^13^C (‰)), δ^15^N (δ^15^N (‰)), soil water content (SWC, g H2O g^-1^ dry soil), and gene copy numbers (gene copies g^-1^ dry soil) in 0-5 (dark brown) and 15-20 (light brown) centimetres soil depth. Data points represent ASV richness (y-axis) and environmental variables or microbial density (x-axis) of individual aggregates. Lines depict calculated regression lines, with shaded areas around them indicating the 95% confident interval for the regressions. We print goodness-of-fit measure (*R*^2^, model accuracy) of the regression models, with asterisks denoting significance levels (*** *P* value≤0.001, ** *P* value≤0.01, * *P* value≤0.05). Highly significant correlations with a *R*^2^>0.1 are highlighted with a bold, coloured border around plots.

## Discussion

In this study, we show that the soil microbiome was structured into distinct subcommunities at the millimetre-scale. Individual 2-millimetre-sized soil aggregates hosted variable bacterial, archaeal, and fungal communities which systematically differed in community composition, richness, and variability compared to bulk soil samples. Furthermore, aggregates and bulk soil samples differed in (proportion of) phyla dominating the soil microbiome. Hence, taxa that dominate communities in individual aggregates and thus might be responsible for certain small-scale biogeochemical processes may not be recognized when analysing large soil volumes. Composition and richness of microbial communities were intimately linked with aggregate-specific physico-chemical variables, as well as the horizontal and vertical sampling locations of aggregates.

The fact that a small fraction of microbial ASVs was exclusively detected in bulk soil samples suggests that some habitats may have been present only in bulk soil samples, and not in 2-millimetre-sized aggregates. This could be, for example, habitats associated with larger pieces of inter-aggregate POM or larger inter-aggregate soil pores. Some fungal taxa seem to not colonise soil aggregates, probably because of their sheer size which might prevent colonisation of aggregate pores. The fraction of ASVs that was exclusively found in aggregates, however, must have also been present in bulk soil samples (as they are a mixture of aggregates), but was not detectable there. As communities of aggregates were, due to their lower soil volume, less diverse than communities found in homogenized parent soil cores, we were able to capture a higher percentage of their total ASVs at a given sequencing depth (Fig. S3). This means that we received a more complete picture of aggregate communities compared to bulk soil communities, including taxa that occur at low relative abundances in individual aggregates. Moreover, we found that any two aggregates taken within a few centimetres differed mostly in their low abundant taxa (Fig. 4), indicating that these rare taxa are also highly heterogeneously distributed at the millimetre-scale. Some of these taxa are thus likely not detectable in bulk soil samples due to their consequently low overall relative abundance. The role of rare taxa in the soil microbiome remains unclear. They could be relics from the past, serve as insurance for changing environmental conditions, or occupy highly specialized niches. Here we show that a certain fraction of them can likely only be observed when sequencing very low soil volumes.

The relationships between microbial communities and physico-chemical variables within aggregates suggest that aggregates provide a stable environment for a sufficient time to allow the development of distinct communities in interaction with their environment. It has been suggested that the relationship between microbial community structure and physico-chemical properties of aggregates is intimate and bidirectional as microorganisms and environment mutually influence each other [4, 5, 60]. Not only the environment selects for taxa with specific traits, but also microorganisms shape their environment [60]. For instance, microbially excreted extrapolymeric substances [22] and sheer presence of fungi [61] have been found to increase wettability of soil. Microbes affect composition and stability of organic matter (OM) [14] by not only transforming but also becoming OM after cell death [62, 63]. It is also likely that fungal and bacterial communities influence each other during the lifetime of individual aggregates. Fungal communities have been shown to shape associated bacterial/archaeal communities in other soil microhabitats, such as mycorrhizal root tips [64]. Consequently, we propose that a similar interdependence between fungal and bacterial communities may also exist within soil aggregates. Examining whether fungal and bacterial/archaeal communities exert an impact on each other’s fine-scale structure in soil is an area that warrants further investigation, which goes beyond the scope of this study.

We can interpret carbon (C) and nitrogen (N) content as proxy for OM content, and δ^13^C and C-to-N ratio as proxy of recycling status of OM as decomposed OM and microbes have a lower C-to-N ratio and are more ^13^C-enriched (more positive δ^13^C) than fresh plant litter [65–68]. Hence, our results demonstrate, for the first time, relationships between microbial community structure and OM content and recycling status at the aggregate-scale. This indicates that soil aggregates may act as functional units of soil organic matter turnover, as has been hypothesised before [69].

Observed shift in microbial community composition between soil depths was accompanied by a systematic difference in C and N content, C-to-N ratio, and δ^13^C between soil depths (Fig. S4). A shift in fungal and bacterial community composition with depth has frequently been found in studies based on bulk soil samples [70, 71]. Moreover, we found that fungal and especially bacterial and archaeal community composition were more similar to each other in the deeper soil layer compared to topsoil (Fig. S5), similar to what has been shown in bulk soil samples [70]. A recent study has demonstrated that diversity and complexity of OM decreased with increasing soil depth [72], which could result in more similar microbial communities. We have not measured diversity and complexity of OM, yet we found indication for reduced variability in elemental composition across aggregates of the lower soil layer compared to the upper one (Fig. S4, Fig. 8). It is also possible that the decrease of community similarity with soil depth is a consequence of decreased microbial richness in the deeper soil layer. Moreover, the weaker relationship between OM content and microbial community composition in 0-5 centimetres soil depth compared to 15-20 centimetres may be due to the masking effect of the overall bigger variability in environmental conditions in the upper soil layer. Conversely, the stronger relationship between OM content and community composition, and less plot-specific bacterial and archaeal community composition in the deeper soil layer could suggest that aggregate-habitats in the deeper soil were overall more similar compared to topsoil.

The weaker relationship between community composition and elemental variables for fungi compared to bacteria and archaea might, for one, be attributed to the presence of saprotrophic fungi who are able to translocate C and nutrients via their hyphae over centimetres to meters [73] and thus can uptake additional C and nutrients outside the investigated aggregate. The mycorrhizal fraction of fungal communities might further weaken the link between OM content and recycling status as mycorrhizal fungi obtain C exclusively from their host plants [74] and thus do not depend on free carbon substrates.

Sampling location, by far, explained the largest proportion of variation in community composition within soil layers. It has been shown that more proximate soil aggregate-parts (parts of the same aggregate) were more similar in community composition than parts of different aggregates [30]. Environmental variation and/or dispersal limitation can underlie the increase in community dissimilarity with increasing horizontal spatial distance [75, 76]. We hypothesize that variability in tree species might have driven observed distance-decay by affecting all microbial communities within a soil core through their residues and exudates equally. In addition to the overall distance-decay exhibited by both bulk and aggregate communities, it was apparent that bacterial and archaeal communities in aggregates were more dissimilar from each other compared to those in bulk soil samples at any given distance. In particular, communities from aggregates taken from the same parent soil core were more dissimilar than bulk soil communities taken from adjacent parent soil cores, and often more similar to aggregate or bulk soil communities of other sampling locations. Together, this exemplifies the high small-scale heterogeneity of aggregate-associated communities and hints to the existence of drivers of community formation operating at the aggregate scale.

The relationship between microbial richness and OM content in the topsoil has a unimodal shape which also has been found for the productivity-diversity relationship of bacteria in soil [77, 78]. Ecological theory proposes that the largest number of taxa co-exists at intermediate substrate concentrations [78, 79]. At low substrate availability, species richness is limited and at high substrate concentrations, few fast-growing taxa might become dominant and outcompete other functional groups [80]. We argue that the positive linear relationship of OM content and microbial richness observed in the deeper soil layer was due to OM content being generally in the lower range.

The unimodal shape of the relationship between bacterial and archaeal density and richness in the topsoil suggests that at highest bacterial and archaeal density fewer ASVs dominate communities than at intermediate density. The missing link between fungal density and richness might be due to fungal gene copy numbers being a rather imperfect measure of fungal density as number of rRNA operon copies per genome varies considerably across fungal taxa [44].

Contrary to our expectation, fungal richness of aggregates was linked to OM content equally strong like bacterial and archaeal richness although organisms differ substantially in size and life strategy. Fungal richness might even have underlain the positive relationships of bacterial and archaeal richness and OM content. The region around hyphae has been denoted to create unique niches for bacteria, offering easily available carbon at the hyphal interface [81] and selecting for certain bacterial taxa [64]. It is likely that high fungal taxonomic richness is accompanied by high functional diversity, potentially resulting in the excretion of a larger range of exudates, which provide more ecological niches and thereby promote the co-existence of a larger number of bacterial and archaeal taxa.

OM diversity and complexity could explain the negative relationship between microbial richness and OM recycling status. Plant litter material is more diverse and complex than decomposed OM [72]. Hence, lower diversity of more recycled OM may constrain co-existence of microbial taxa due to reduced resource differentiation. Also, potentially enhanced mineral stabilisation of further decomposed OM may limit taxa co-existence by reducing substrate availability.

Increased soil moisture might have promoted bacterial and archaeal richness due to enhanced bacterial and archaeal movement and solute transport resulting from improved connectivity of the soil pore space [5, 7]. Explaining the observed positive relationship of fungal richness and soil moisture is more difficult as fungi do not depend on a continuous water-film to reach substrates and new habitats as they are predominantly aerobic and can gap air-filled pores [61, 82]. Probably, OM content has underlain the positive relationship between microbial richness and soil moisture as presence of OM has been shown to increase water content and water holding capacity of soils [83, 84].

The findings of our study are derived from samples originating from a specific temperate forest soil. They may not be transferrable to other soil types and ecosystems as the status and dynamics of aggregates are influenced by various factors such as the type of vegetation, edaphic parameters, soil texture, organic matter concentration and abiotic conditions. To improve our understanding of the general patterns of small-scale organisation of the soil microbiome, further research examining aggregate-scale communities and their immediate environments in different ecosystems would be beneficial. When studying soil aggregates ex-situ careful and rapid handling is crucial to avoid changes in soil water content and minimize changes in surface environmental conditions due to air exposure, which can potentially distort findings.

## Conclusion

We conclude that the soil microbiome is a metacommunity comprising variable subcommunities at the small-scale, which collectively contribute to the vast richness of the soil microbiome. The variability in subcommunity composition and richness is closely related to the heterogeneous distribution of OM with different recycling status and water throughout the soil matrix, as well as the spatial distance between communities. Our study emphasizes the necessity of studying small, coherent soil units to enhance our understanding of the controls of the structure of within-itself connected microbial communities and, ultimately, community-mediated biogeochemical processes in soils.

## Supporting information

Supplemental Material textsummary

## Acknowledgments

We thank Barbara Kitzler for helpful support in selecting and accessing the research site, Lilian Kaufmann for help in the field, and Julia Mor for help in the lab, Margarete Watzka for measuring elemental composition of samples, Syrie Hermans for the helpful, detailed exchange (including R scripts) about variation partitioning. This work was funded by the European Research Council under the European Union’s Horizon 2020 research and innovation programme (grant agreement No 819446 to CK).

## Data availability

Raw 16S rRNA genes and ITS2 amplicon sequencing data have been deposited in the NCBI Sequence Read Archive under the BioProject accession number PRJNA1090291. All additional data analysed in this manuscript are available on the Zenodo repository (https://doi.org/10.5281/zenodo.10814159).

## Author contributions

CK conceptualised the study. ES, KG, LA, and CK performed field sampling. ES, CR, KJ, and LA performed sample processing and/or analyses in the lab. ES coordinated field and lab work. PP advised and carried out sequencing efforts, HS advised and supported the measurement of fungal and bacterial gene copy numbers by droplet digital PCR. ES conducted data analysis, with contributions from SD. ES wrote the original manuscript draft supported by CK. All authors contributed to manuscript revisions.

