## Supplemental Material for "Distinct microbial communities are linked to organic matter properties in millimetre-sized soil aggregates"

Supplementary Graphs and Tables

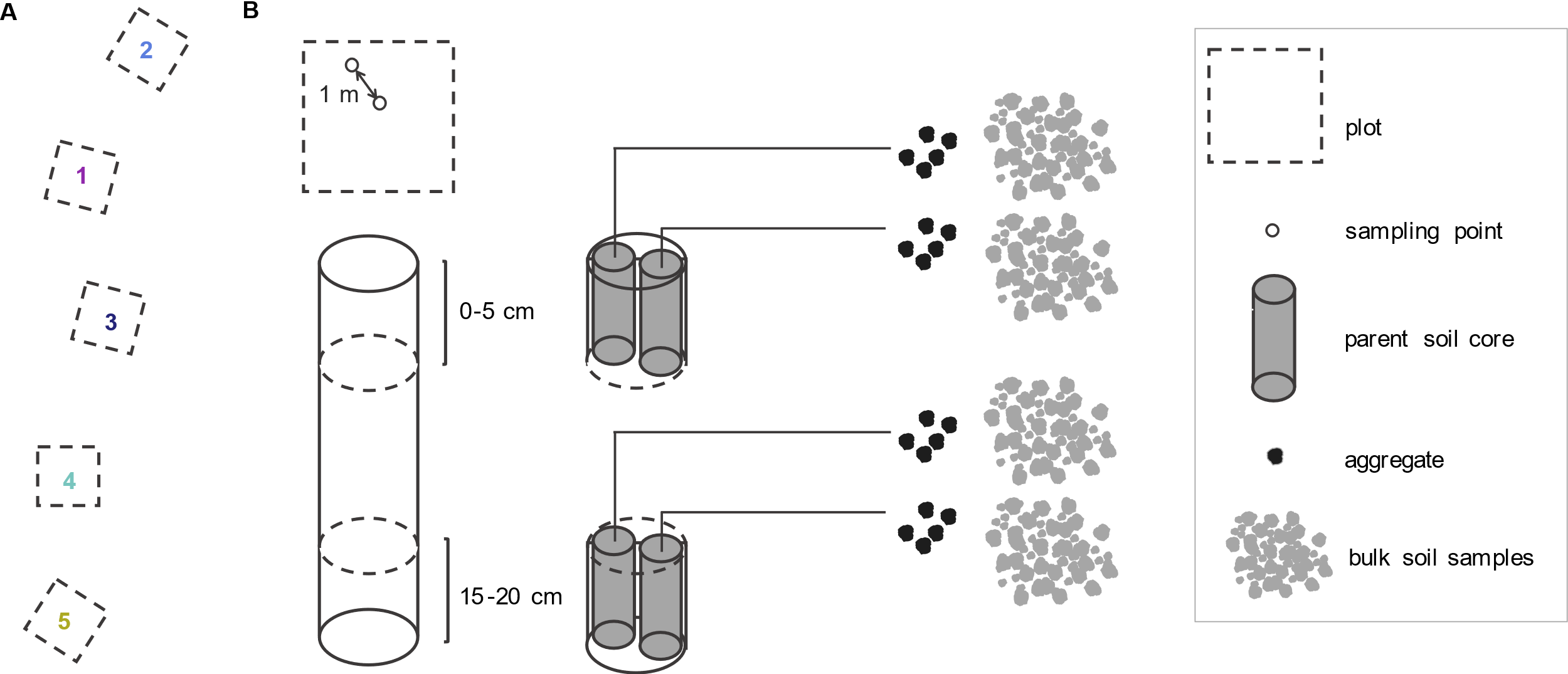

**Fig. S1 Sampling scheme.** (**A**) Spatial arrangement of the five 5x5 meter plots at the forest site. (**B**) Schematic representation of the sampling procedure. Within each plot, we identified two sampling points approximately one meter apart. At each sampling point, we sampled two soil cores (8 cm diameter, 5 cm height), one from 0-5 centimetres and another from 15-20 centimetres. From each core, we subsampled two smaller soil cores (*parent soil cores,* 35 cm^3^ volume, 3 cm diameter, 5 cm height). Subsequently, we hand-picked five individual approximately two-millimetre-sized aggregates from each parent soil core and sieved the remaining soil to two millimetres, producing bulk soil samples.

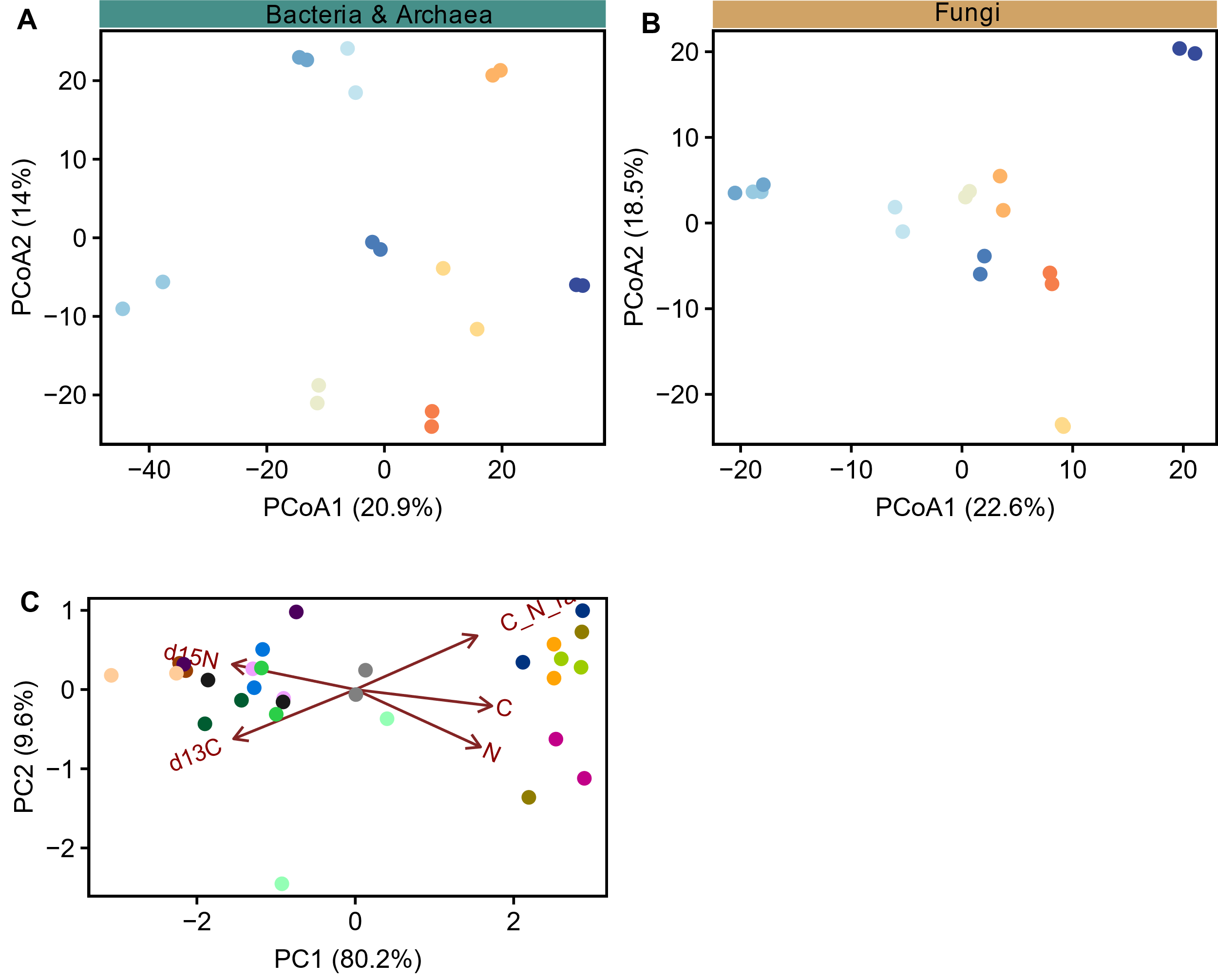

**Fig. S2** **Homogenisation of individual aggregates produced two subsamples with similar community composition and elemental composition.** PCoA plots illustrate (**A**) bacterial and archaeal and (**B**) fungal community similarity (Aitchison distances) of aggregate halves from nine individually homogenised 2-millimetre-sized aggregates. (**C**) The principal component analysis (PCA) plot shows the similarity of elemental composition between the two subsamples from 15 individually homogenised 2-millimetre-sized aggregates. PCA was performed on scaled and centered elemental variables (C and N concentration, C:N ratio, δ^13^C and δ^15^N). Red arrows indicate elemental variables. PCoA plots were generated for aggregates sampled and sequenced alongside aggregates analysed in the main part of this manuscript. Aggregates for comparing elemental composition of aggregate subsamples were sampled in Spring 2021 at the same forest site in Klausen-Leopoldsdorf. All aggregates were freeze-dried (for 48 h) and carefully homogenised by hand using a balltool. Subsamples for assessing elemental composition similarity were analysed with an EA-IRMS. Colours indicate aggregate membership.

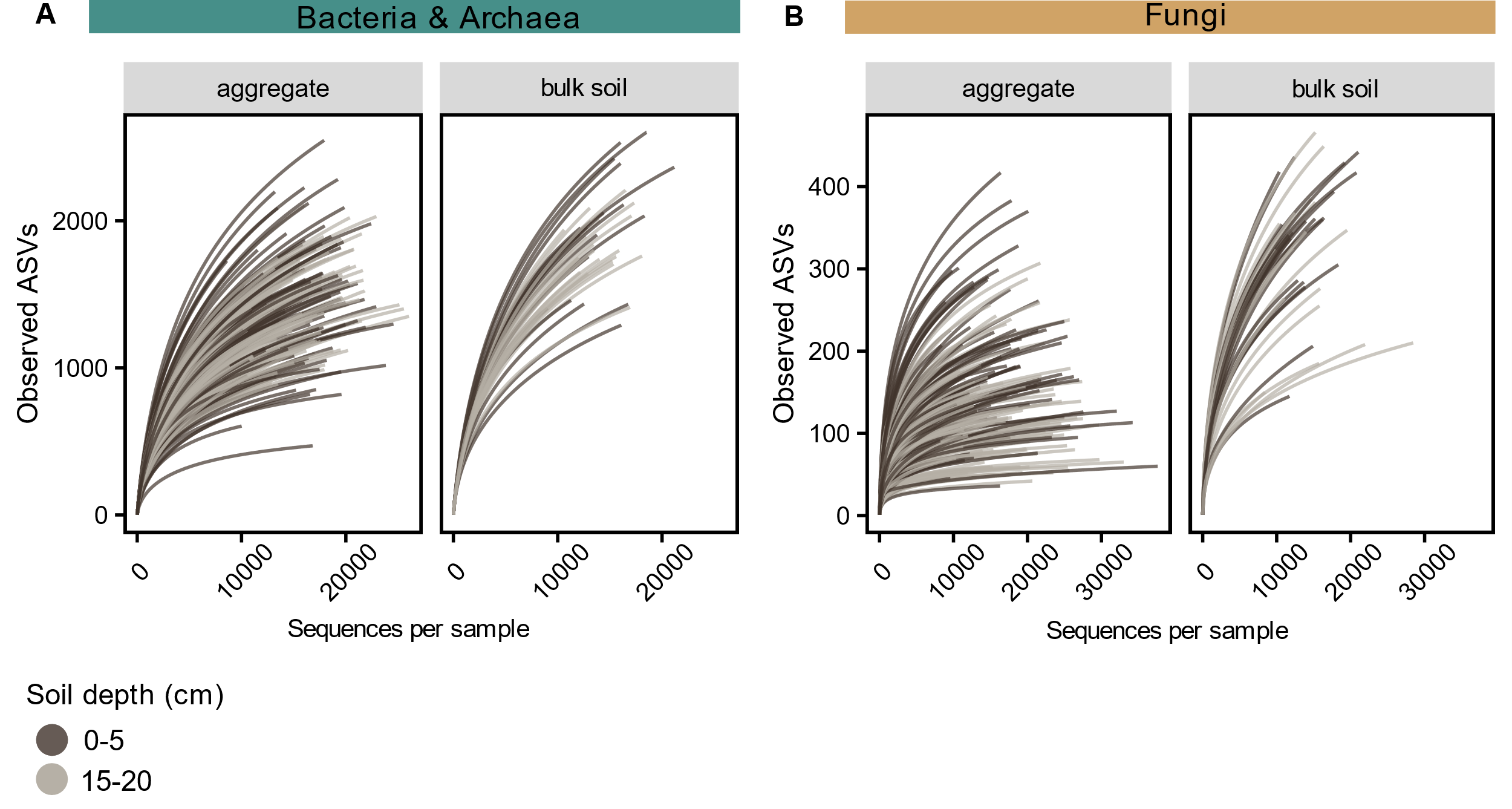

**Fig. S3** **Bacterial and archaeal, and fungal communities within aggregates were captured more completely compared to those in bulk soil samples.** Rarefaction curves illustrate how complete we captured (**A**) bacterial and archaeal and (**B**) fungal communities within individual aggregates (left) and bulk soil samples (right) of 0-5 (dark brown) and 15-20 (light brown) centimetres soil depth, relative to the number of sequences. Curves approximating a plateau indicate a capture of all ASVs present in a community.

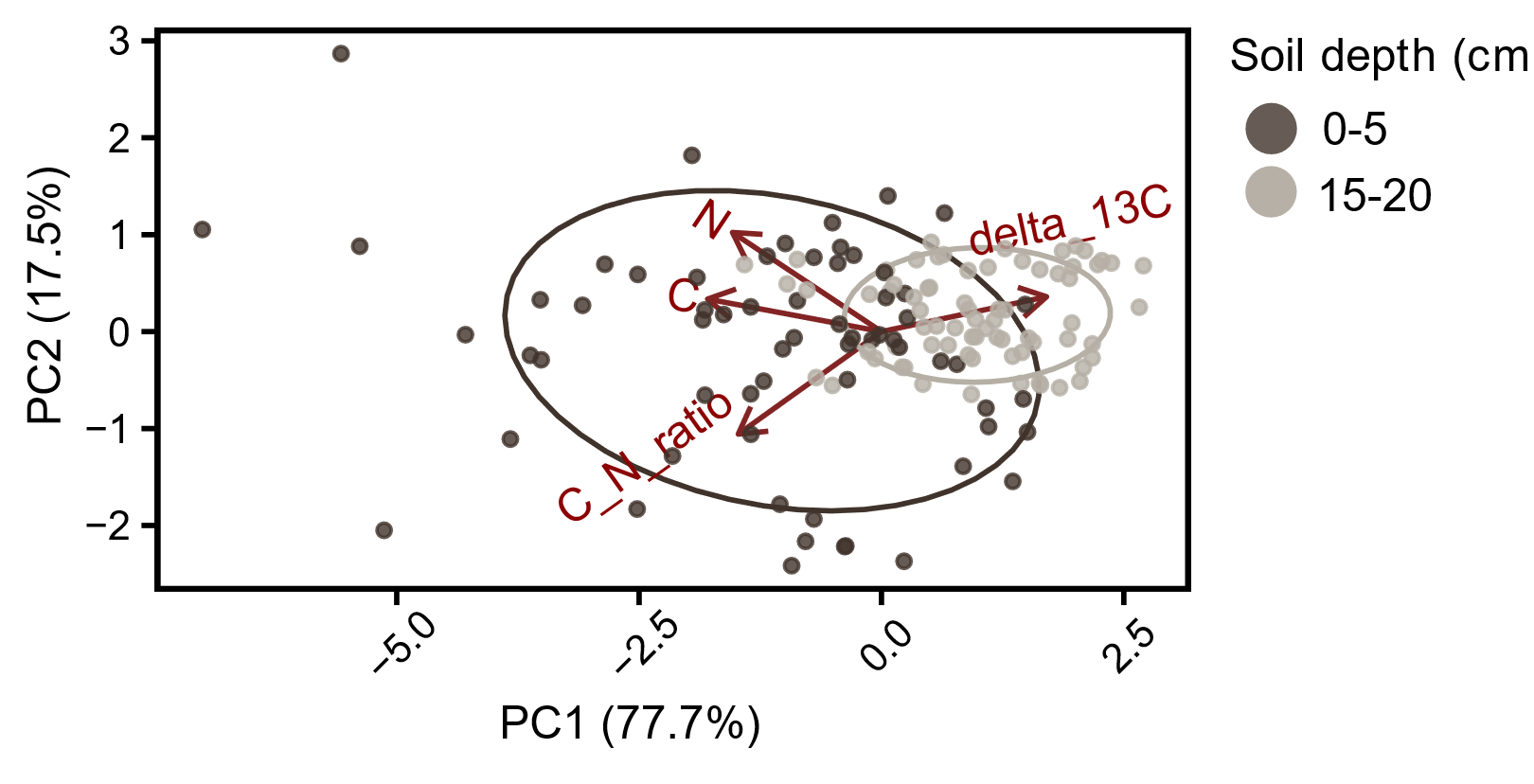

**Fig. S4** **Aggregates systematically differed between topsoil and lower soil layer regarding their elemental composition, with elemental composition of aggregates of topsoil being more variable.** The principal component analysis (PCA) plot visualises the similarity of elemental composition among and between aggregates of 0-5 (dark brown) and 15-20 (light brown) centimetres soil depth. PCA was conducted on scaled and centered elemental variables (C and N concentration, C:N ratio and δ^13^C). Red arrows indicate elemental variables.

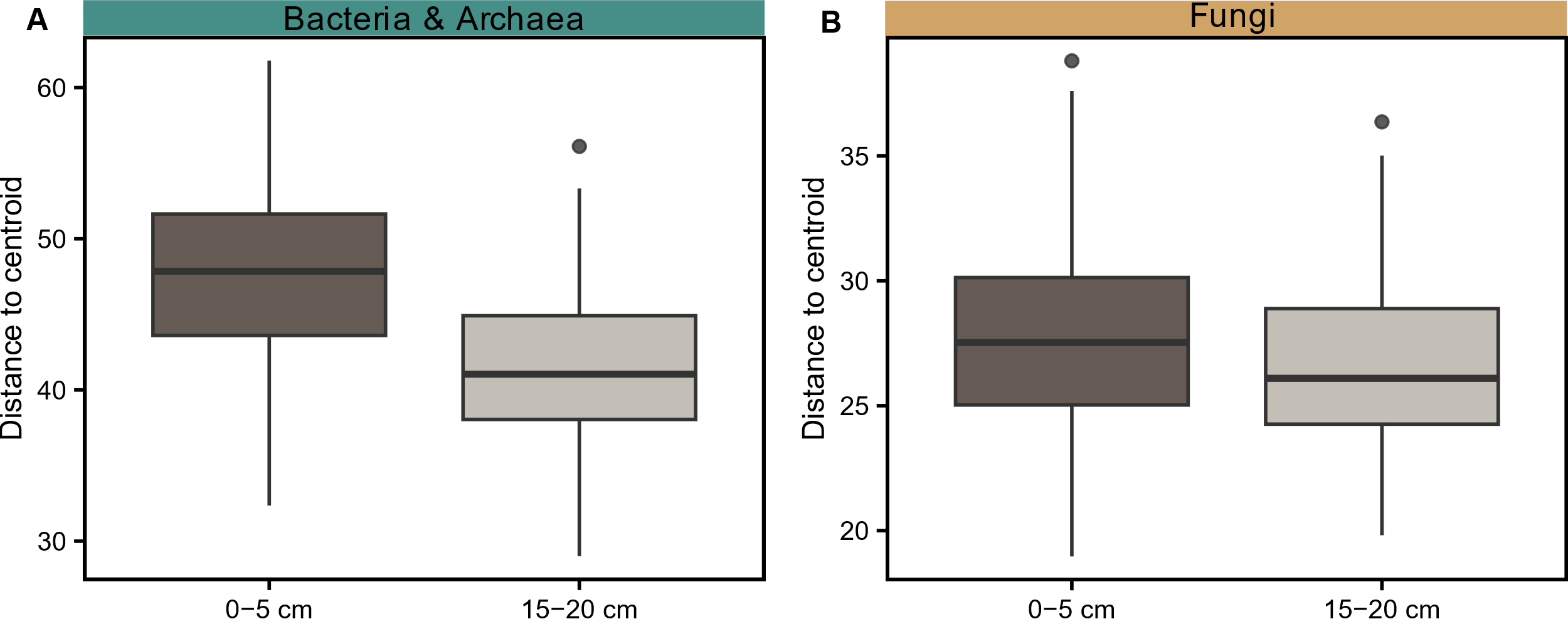

**Fig. S5** **Microbial aggregate communities in the topsoil were more variable regarding community composition than communities in the deeper soil layer.** Boxplots visualise the average distance of an aggregate community from the centroid in 0-5 (dark brown) and 15-20 (light brown) centimetres soil depth based on (**A**) bacterial and archaeal, and (**B**) fungal community dissimilarity (Aitchison distances) in PCoA plots (Fig. 3A, 3B). The centroid is calculated as the gravitational center of aggregates within the same soil layer.

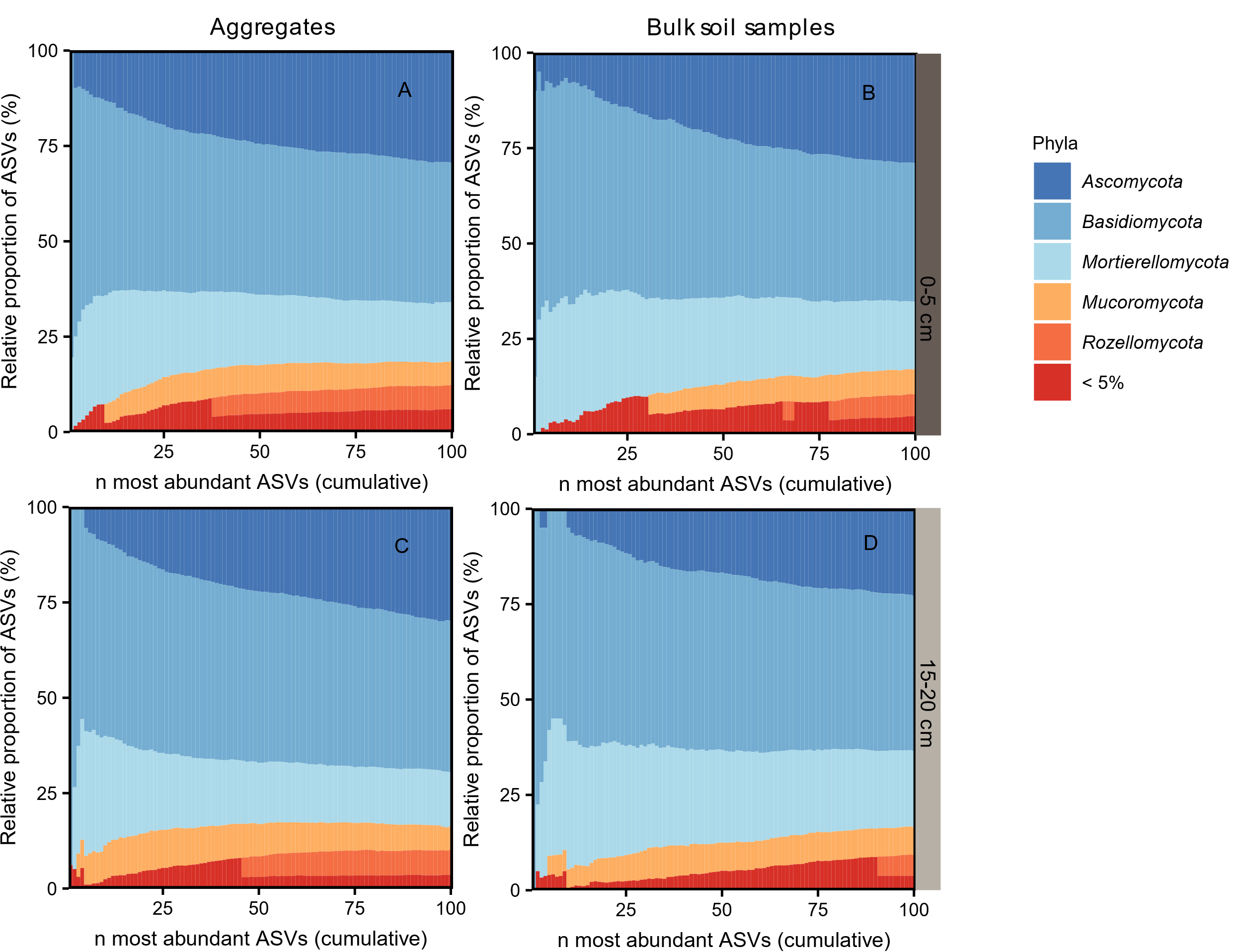

**Fig. S6 Aggregates and bulk soil samples differed slightly in phylum composition of their most abundant fungal taxa.** Stacked bar plots illustrate the phylum composition of 100 abundance categories for (**A, C**) aggregates (0-5 cm: n=92,15-20 cm: n=98) and (**B, D**) bulk soil samples (0-5 cm: n=20, 15-20 cm: n=20) in 0-5 and 15-20 centimetres soil depth. Each stacked bar represents one abundance category (x-axis) and displays the proportions of ASVs belonging to different phyla (y-axis). Abundance categories comprise different numbers of ASVs. Which ASVs are included depends on their ranks which were assigned to ASVs in each sample individually based on their relative abundance. The number of abundance category (1 to 100) indicates how many of the most abundant ASVs (per sample) were included. For instance, abundance category 25 encompasses the 25 most abundant ASVs of all samples. Phyla comprising more than 5% of all ASVs per abundance category are depicted in different colours, while phyla constituting less than 5% of ASVs are grouped into the phylum category “< 5 %”. The analysis is based on non-rarefied datasets.

**Table S1 PCR cycling conditions used for amplicon sequencing of 16S rRNA genes and ITS2 regions.** PCR reactions (25 µl) comprised 1x DreamTaq Green PCR master mix, 0.1 µg µl-1 BSA, 0.25 (for 16S rRNA) or 0.5 (for ITS2) µmol l-1 of each primer and 2 µl of DNA template.

|  | **Temperature (°C)** | **Time** |
| --- | --- | --- |
| **ITS2,** ISOF-T/ITS4 | 94 | 4 min |
|  | 94 | 45 sec (35 cycles) |
|  | 48 | 45 sec |
|  | 72 | 1 min |
|  | 72 | 10 min |
| **ITS2,** Nested PCR (gITS7/ITS4) | 94 | 4 min |
|  | 94 | 30 sec (20 cycles) |
|  | 53 | 30 sec |
|  | 72 | 30 min |
|  | 72 | 10 min |
| **16S** V4 (515F/806R) | 94 | 3 min |
|  | 94 | 45 sec (30 cycles) |
|  | 52 | 60 sec |
|  | 72 | 60 min |
|  | 72 | 10 min |

**Table S2 Primers and PCR cycling conditions for the thermocycler used for Digital Droplet PCR to amplify 16S rRNA genes and ITS1 regions within droplets.** PCR reactions (22 µl) consisted of 11 µl QX200 ddPCR EvaGreen Supermix (1x), 0.2 µl of forward primer (0.1 µM) and 0.2 µl of reverse primer (0.1 µM), 8.6 µl DNase-free water and 2 µL DNA template. Adjustments were made to the ratio between DNA template and reagents when additional DNA template was required to achieve an optimal DNA concentration of 0.1 (for 16S rRNA) and 0.25 (for ITS1) ng/µl for PCR reactions.

| **16S rRNA genes** (515F [1] and 806R [2]) | **ITS1 regions** (ITS1F & ITS2 [3]) |
| --- | --- |
| 95°C for 5 min | 95°C for 5 min |
| 5 cycles of 95 °C for 30 s and 57°C for 2.5 min (−1 °C each step) | 5 cycles of 95°C for 30 s and 60°C for 2.5 min (−1°C each step) |
| 35 cycles of 95°C for 30 s and 52°C for 2.5 min | 35 cycles of 95°C for 30 s and 55°C for 2.5 min |
| 4°C for 5 min | 4°C for 5 min |
| 90°C for 5 min | 90°C for 5 min |
| 10°C hold | 10°C hold |
| Storage at 4°C (min 1 h) before droplet reading | Storage at 4°C (min 1 h) before droplet reading |

**Table S3 Number of pairwise comparisons underlying individual datapoints in Fig. 3 differs.** Each number gives the number of comparisons of a certain spatial distance category: samples from the same parent soil core (spsc), adjacent parent soil cores (apsc), from approximately one-meter apart parent soil cores (1), 10-20 meters (10-20), 20-30 meters (20-30), 30-40 meters (30-40), 40-50 meters (40-50), 10-20 meters (50-60) apart parent soil cores.

|  | Spatial distance categories | | | | | | | |
| --- | --- | --- | --- | --- | --- | --- | --- | --- |
|  | **spsc** | **apsc** | **1** | **10-20** | **20-30** | **30-40** | **40-50** | **50-60** |
| **0-5 cm** | | | | | | | | |
| Bacteria & Archaea | | | | | | | | |
| aggregates | 152 | 188 | 338 | 831 | 460 | 666 | 506 | 180 |
| bulk soil samples |  | 9 | 16 | 40 | 25 | 32 | 24 | 8 |
| Fungi | | | | | | | | |
| aggregates | 152 | 188 | 338 | 831 | 460 | 666 | 506 | 180 |
| bulk soil samples |  | 9 | 16 | 40 | 24 | 32 | 24 | 8 |
| **15-20 cm** | | | | | | | | |
| Bacteria & Archaea | | | | | | | | |
| aggregates | 172 | 215 | 380 | 950 | 580 | 761 | 570 | 200 |
| bulk soil samples |  | 9 | 16 | 40 | 24 | 32 | 24 | 8 |
| Fungi | | | | | | | | |
| aggregates | 172 | 215 | 380 | 950 | 580 | 761 | 570 | 200 |
| bulk soil samples |  | 9 | 16 | 40 | 24 | 32 | 24 | 8 |

**Supplementary Material and Methods**

**Field soil core transport and parent soil core sampling**

We transported intact soil cores (diameter=8 cm, height=5 cm) from the field to the lab. We ensured intactness of field soil cores by transporting them in cut plastic bottles which were of the same diameter and height as soil cores. Soil cores in plastic bottles were kept separately in closed plastic bags to avoid drying of soil. In the lab, we carefully subsampled two smaller soil cores (diameter=3 cm, height=5 cm) from each field soil core, which we call *parent soil cores*. We stored sampled soil cores in separate plastic bags to avoid air drying of soil.

**Aggregate sampling and weighing**

We sampled 5 approximately 2-millimetre-sized soil aggregates from each parent soil core. We took special care to ensure as little drying of soil aggregates during sampling and weighing. This was guaranteed by our sampling setup and a handling duration of a few minutes.

In detail, we set up a petri dish (with an approximate diameter of 8 cm) containing a ring of moist filter paper which we had wetted with milliQ water to avoid drying of aggregates while in the petri dish. Underneath the petri dish, we placed a piece of millimetre paper to select aggregates with an approximate diameter of two millimetres. We transferred a small amount of soil (less than a teaspoon) in the centre of the petri dish, making sure soil did not touch the moist filter paper. We selected one aggregate after another. Each selected aggregate was placed in a fresh, tared metal tin with the help of a fine tweezer and weighed on a fine mass balance (Sartorius Cubis MSE 36P, precision=0.001 mg) to determine its fresh weight. After weighing, the aggregate was transferred into a safe-lock tube which we closed immediately. The whole procedure of picking an aggregate over weighing it until closing the tube containing the aggregate took only few minutes. The petri dish was closed with a lid between picking individual aggregates to avoid drying of soil remaining in the petri dish. Whenever soil in the petri dish appeared to have dried, we discarded it and replaced it with a new teaspoon of soil.

Intact aggregates in tubes were stored at -80°C. Individual, still intact aggregates were weighed again after they had been freeze-dried for 48 hours to determine their dry weight.
