## Supplemental Material textsummary for "Distinct microbial communities are linked to organic matter properties in millimetre-sized soil aggregates"

Text Summary, Supplementary Material

The Supplementary Material file (.docx) contains 6 supplementary figures and 3 supplementary tables. They either provide relevant additional information for the Material and Methods section or support claims in the Discussion section of our paper.

Furthermore, we include a Supplementary Material and Methods section describing soil sampling in the field, core transport to the lab and aggregate sampling and handling in detail.
